# The endoderm cell trajectory of urochordate *Styela clava* reveals the dual developmental origin and evolution of digestive tract

**DOI:** 10.64898/2026.01.25.701569

**Authors:** Yonghang Ge, Wei Zhang, Penghui Liu, Jianqing Bi, Haiyan Yu, Bo Dong, Jiankai Wei

## Abstract

The digestive system exhibits extensive diversity in developmental mechanisms and morphology across metazoans, yet the evolutionary origins underlying its organ differentiation remain unclear. Here, single-cell RNA sequencing was employed to investigate endodermal cell lineage specification during metamorphosis in the urochordate *Styela clava*, a newly established model for chordate evolution. By profiling 26,099 cells across five stages, we identified 21 major cell clusters and reconstructed the endodermal differentiation trajectories. Our analysis reveals two larval endodermal progenitor populations with distinct differentiation potentials. Pseudotime and RNA velocity analyses indicate that these progenitors give rise to stomach and intestinal lineages, respectively. Cross-species comparisons reveal putative homologous relationships between ascidian endodermal lineages and mouse definitive and visceral endodermal lineages, suggesting dual origins of digestive tract in chordates. We also identified conserved TGF-β and FGF regulatory programs in digestive organ patterning and highlight earlier fate restriction of stomach and intestinal progenitors in ascidians compared to vertebrates. These findings provide insights into how chordate digestive organs evolved from ancestral endodermal patterning programs.

## Background

The digestive system, comprised of digestive tract and accessory organs, plays a central role in nutrient acquisition and processing across animals. While exhibiting morphological diversity depending on habitats and feeding strategies, digestive systems universally share conserved functional processes including ingestion, digestion, absorption, and elimination.^1^

Developmentally, the digestive tract and accessory organs are primarily originated from endodermal derivatives. However, the origins and formation patterns of these organs exhibit notable differences between invertebrates and vertebrates.^2^ For example, in *Drosophila melanogaster*, the midgut derives from endoderm while foregut and hindgut originate from ectoderm.^3^ In *Caenorhabditis elegans*, both ectodermal and mesodermal cells contribute to its foregut.^4^ By contrast, in vertebrates especially mammals, digestive tracts are entirely endoderm-derived, with fore-, mid- and hindgut all arising from this germ layer.

One reason for this distinction lies in the profound differences of behaviors and origins of endodermal cells during early development. The endoderm cells of invertebrate remain internalized after gastrulation, whereas vertebrate endoderm undergoes dynamic morphogenetic movements.^2,5^ Moreover, mammalian endoderm development displays additional complexity through its dual origin. The definitive endoderm (DE) arises during gastrulation from epiblast cells, while extra-embryonic (primitive endoderm) forms earlier from the inner cell mass. These two endodermal lineages collectively give rise to the embryonic gut endoderm and resulting gut tube.^6^

Furthermore, there is also distinct formation patterns during organogenesis of digestive organs between invertebrates and vertebrates. For example, in vertebrates, the gut tube first becomes regionalized into foregut, midgut, and hindgut. As development proceeds, The foregut gives rise to the esophagus, trachea, stomach, lungs, thyroid, liver, biliary system, and pancreas; whereas the midgut forms the small intestine and the hindgut forms the large intestine.^7^ But unlike the highly compartmentalized and organ-specific digestive systems of vertebrates, invertebrate gut structures are generally less complex and exhibit more diffuse regional specialization.^8–11^ These developmental differences underscore distinct cellular lineages between invertebrate and vertebrate digestive systems, which has hindered our understanding of how digestive tracts originated and evolved from invertebrates to vertebrates.

Urochordates, as the closest living relatives of vertebrates, offer a valuable model for exploring the developmental and evolutionary origins of the vertebrate organs.^12^ Their unduplicated genomes and conserved developmental pathways make them especially informative for reconstructing ancestral chordate traits.^13^ They also provide insights into the evolutionary origins of cell types in vertebrates.^14^ Recent advances in comparative genomics using *Styela clava* have facilitated cross-species analyses of gene regulatory networks involved in organ evolution.^15,16^

In ascidians, the digestive system developed during metamorphosis and composed of pharynx, esophagus, stomach, intestine and pyloric gland, lacking complex accessory organs and glands.^17^ Although the structure of ascidian digestive system is relatively simple, the basic function in nutrition processing is similar to vertebrates.^18,19^ Previous studies have shown that the genes encoding enzymes involved in digestion, absorption, and immune function in ascidian are very similar to those of vertebrates.^20–22^ Meanwhile, the expression patterns of some transcription factors are also conserved between ascidians and vertebrates.^1,20^ However, the cellular heterogeneity and lineage dynamics of the urochordate digestive tract remain largely unexplored especially at single-cell resolution.

To investigate the evolution and diversification of digestive systems, it is essential to integrate not only morphological data but also information on cellular lineage and molecular regulation. While vertebrate models, particularly mammals, have provided extensive insights into digestive organ morphogenesis and function, the evolutionary history that gave rise to these systems are less well characterized. Here, we employ single-cell RNA sequencing to map the transcriptional landscapes and developmental trajectories of digestive tract cells in a urochordate model. Our analysis reveals conserved regulatory programs but divergent cell specification strategies underlying digestive tract formation, and provides new insights into the cell lineage evolution of digestive tract across chordates.

## Results

### The single-cell atlas of ascidian *S. clava* across metamorphosis

In ascidian, endodermal organogenesis is mainly occurred during metamorphosis. In order to trace the crucial developmental events, we performed single cell RNA-sequencing (scRNA-seq) using samples from five developmental stages including hatched swimming larva (hsl), tail regressing larva (trl), tail regressed larva (trj), metamorphic juvenile (mj), and juvenile (j) stages. These stages span the main metamorphotic processes of *S. clava* (Figure 1A). After quality control, a total of 26,099 cells were acquired, with an average expression of 11,413 genes per stage (Figure S1). The major cell types were identified (11, 11, 15, 17, and 17 cell types at hsl, trl, trj, mj, and j stage, respectively) based on cluster-specific marker gene expressions (Figure S2, Table S1). The Uniform Manifold Approximation and Projection (UMAP) plots displayed the cell type composition at each stage (Figure 1B). And the cell type dynamic changes were observed during metamorphosis. For example, notochord cells disappeared at trj and mj stages, while endostyle and digestive tract cells appeared at mj and j stages (Figure 1B). To compare the cell type consistency, the cell type similarity heat map of adjacent stages was acquired based on the gene expression profiles. The results showed a higher one-to-one consistency between hsl and trl stage, and between trl and trj stage (Figure 1C). The integrated UMAP of sc-RNA seq data also revealed that the cell types from stages before metamorphosis (hsl, trl and trj stages) were distinctly separated from the ones after metamorphosis (mj and j stages) (Figure 1D).

**Figure 1.**
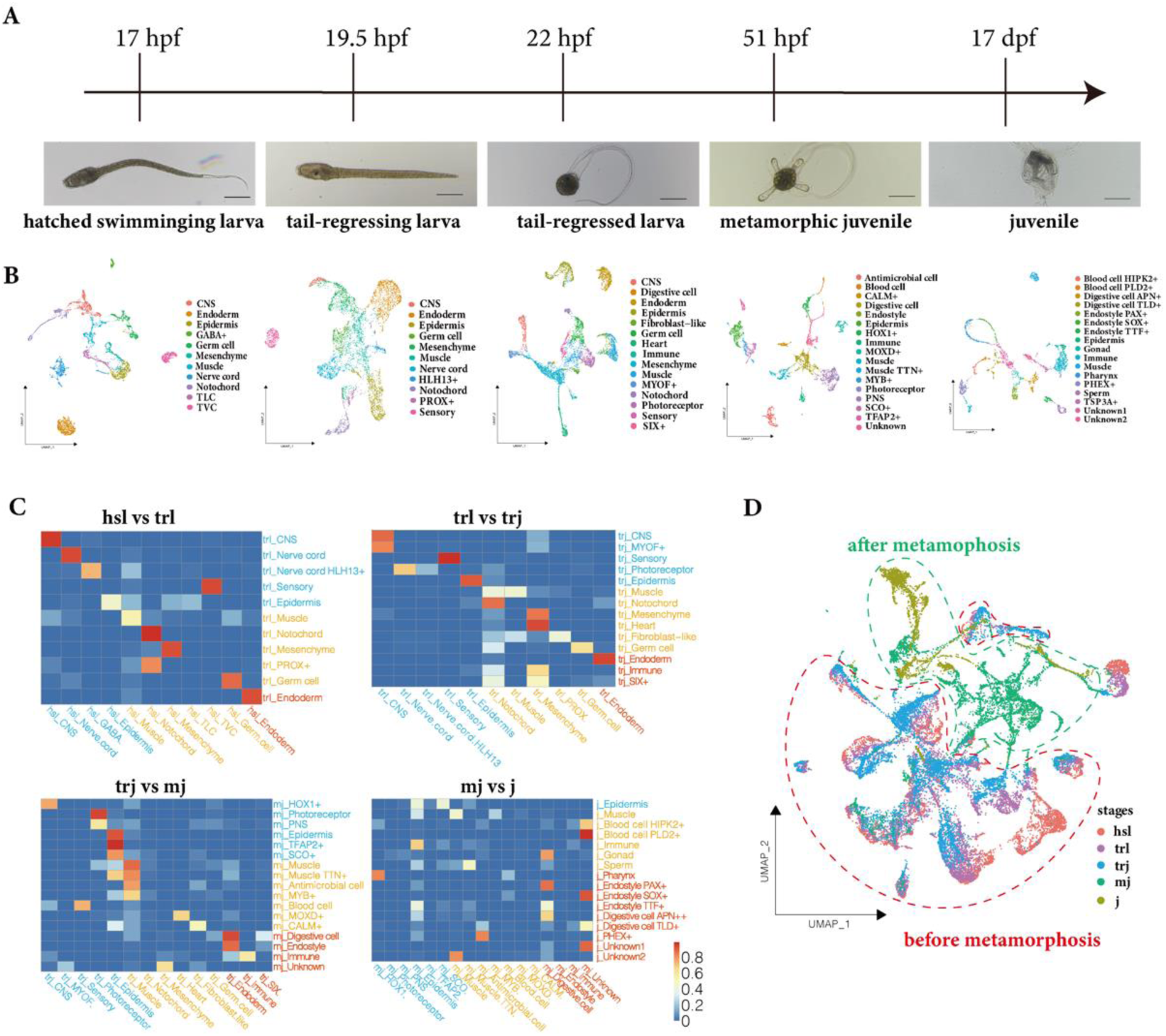
Identification of major cell types across five developmental stages in *S. clava*. **(A)** Representative sampling stages and corresponding larval and juvenile morphologies. (**B**) Cell type annotations derived from scRNA-seq data, aligned with the respective developmental stages. Hpf, hours post-fertilization; Dpf, days post-fertilization. (**C**) Inferred corresponding relationships between cell types across successive stages. Each cell type at a later stage (y-axis) is matched to its most similar cell type from the previous stage (x-axis) based on transcriptomic similarity calculated via K-nearest neighbor (KNN) analysis. A similarity score > 0.7 is used as the criterion for inferring the corresponding relationship. Cell types are color-coded by their germ layer of origin: ectoderm (blue), mesoderm (yellow), and endoderm (red). (**D**) UMAP plot integrating scRNA-seq data from all five stages, with cells colored by developmental stage, illustrating the cellular state transitions before and after metamorphosis. Cells from pre-metamorphic stages are outlined by red dashed lines, while post-metamorphic cells are enclosed by green dashed lines.

### Increased proportion of endoderm-derived cells after *S. clava* metamorphosis

To gain a global single-cell atlas, we integrated the data of all stages and obtained a total of 72 cell-types (Figure 2A). According to the origin of germ layers, we divided them into ectoderm derived cells (including epidermal cells, central nervous system (CNS), peripheral nervous system (PNS), gamma-aminobutyric acid cells, nerve cord cells, sensory cells and photoreceptor cells), mesoderm derived cells (including mesenchyme cells, trunk lateral cells (TLCs), trunk ventral cells (TVCs), notochord cells, muscle cells (body wall muscle, heart muscle), blood cells, and germ cells) and endoderm derived cells (including endostyle cells, digestive cells, and pharynx cells). The expressions of germ layer restricted maker genes showed that endodermal makers, such as *Nkx2-1, Nkx2-6, A1CF* and *THA* were expressed in endoderm-derived cell types, and mesodermal markers (*Noto8, FGF18, KDM8, TPM*, *TNNC*) and ectodermal markers (*Ces, NRCAM, SCO, SSPO*) were also expressed in mesoderm and ectoderm derived cell-types, respectively (Figure 2B, Figure S2A, Table S1), which validated the cell type annotation and classification. In order to display the cell dynamics of each germ layer during metamorphosis, the cells were labeled based on their germ layer origins before (hsl, trl and trj stages) and after metamorphosis (mj and j stages) on UAMP (Figure 2C). The result not only showed a distinct cell composition but also an increasing of proportion of endodermal derived cells after metamorphosis. The ectodermal and mesodermal derived cell types occupied the majority before metamorphosis, while after metamorphosis the proportion of endodermal derived cells were increased from trj to j stage (Figure 2D).

**Figure 2.**
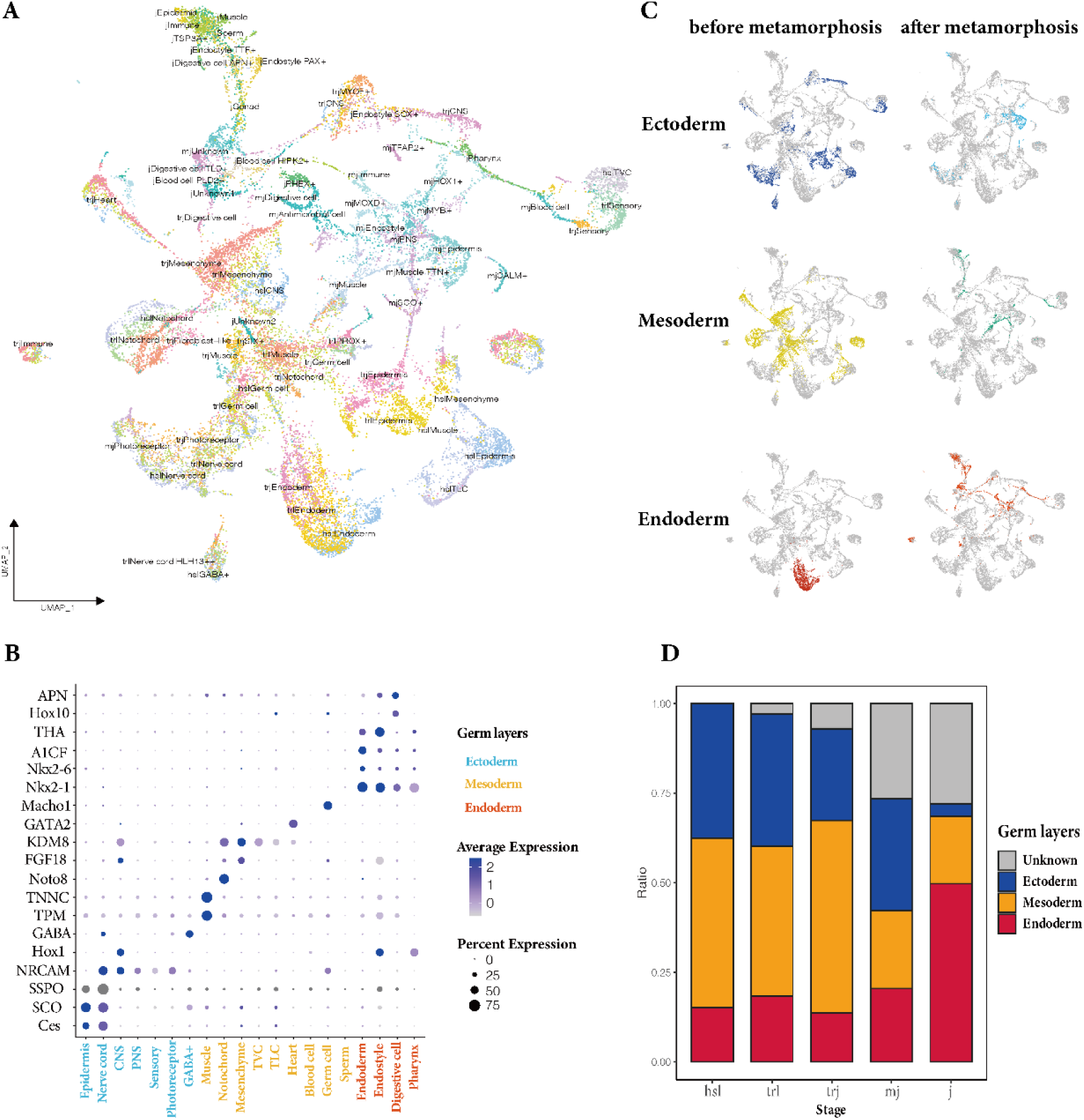
Germ layer-based cell type composition and dynamics during metamorphosis of *S. clava*. (**A**) UMAP plot of all 26,099 cells integrated across five developmental stages, showing the distribution of cell types. Cluster labels represent a combination of developmental stage and cell type identity. (**B**) Dot plot displaying the expression of representative germ layer-specific marker genes. Dot color indicates the average expression level within each cluster, and dot size reflects the proportion of cells expressing the gene. (**C**) UMAP plot highlighting cells derived from the three germ layers, showing the distribution of ectoderm-, mesoderm-, and endoderm-derived populations before and after metamorphosis. (**D**) Bar plot illustrating the proportional changes in cell populations derived from each germ layer across the five developmental stages.

### Two distinct endodermal populations with different differentiation potential

To investigate the differentiation of endoderm-derived cells during metamorphosis, we selected 1,815 cells from trj stage (before metamorphosis) and mj stage (after metamorphosis) with expression of endoderm-specific marker genes (Figure S2B). The sub-cluster analysis identified 12 cell types, including four cell types of endoderm cells from trj stage, and three cell types of stomach progenitors, three cell types of intestine progenitors, endostyle cells, immune cells from mj stage, representing the endodermal cell lineage during metamorphosis of *S. clava* (Figure 3A). Then, the differentiation potential levels of each cell type were predicted using CytoTRACE analysis, with higher scores indicating less differentiated, progenitor-like cells and lower scores corresponding to highly differentiated, lineage-committed cells (Figure 3B). The results showed that cell types with different differentiation potentials (ranging from high to low) are observed at both stages (trj stage and mj stage) (Figure 3C and 3D). The differentiation potential level of four endoderm cell types at trj stage showed significant differences.

**Figure 3.**
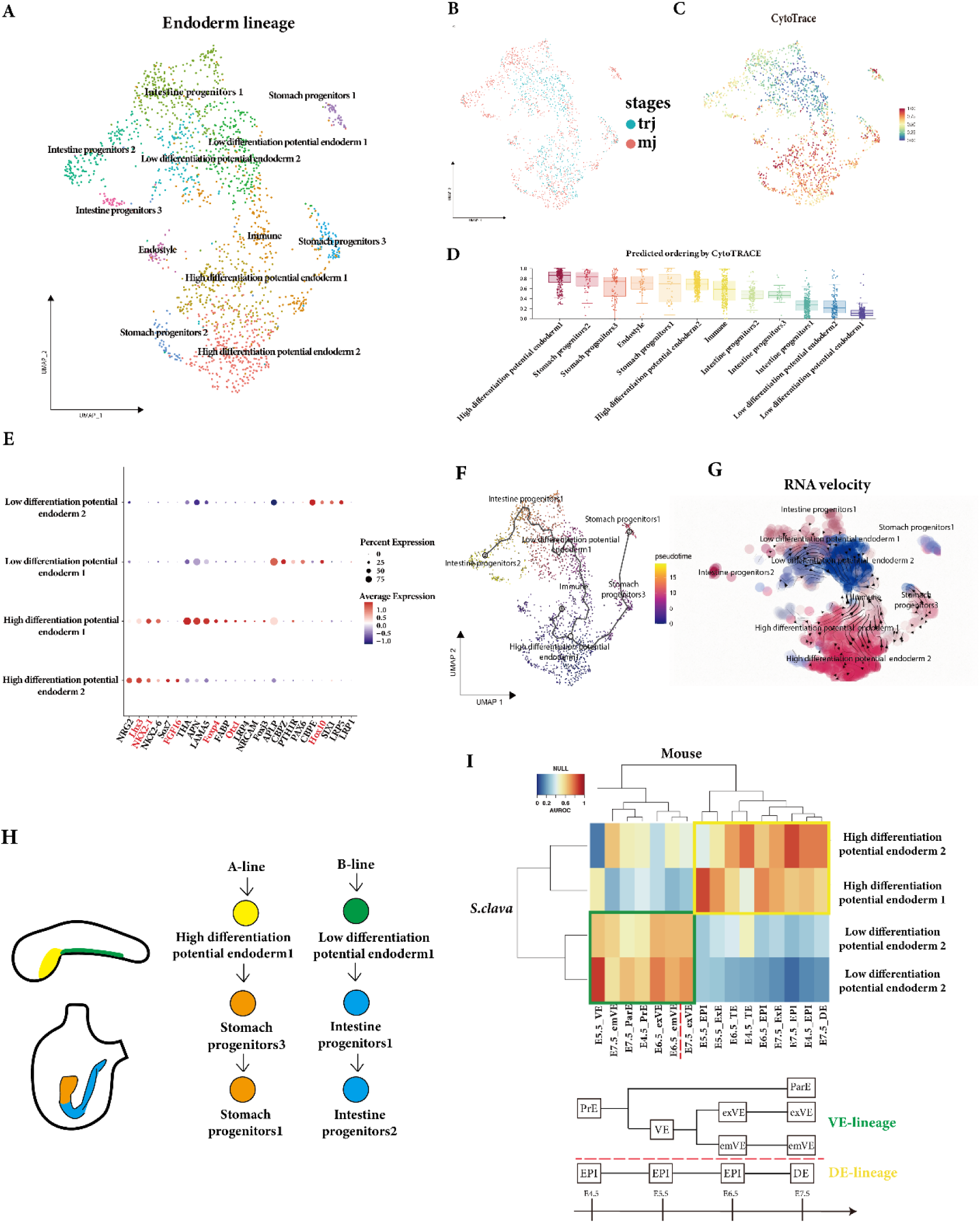
Developmental trajectories of endoderm lineage cells. (**A**) UMAP plot showing sub-clustering of endodermal lineage cells and their distribution during metamorphosis. (**B**) Box plot summarizing the predicted differentiation levels of endodermal lineage cells based on CytoTRACE scores. (**C**) UMAP overlaid with CytoTRACE differentiation scores of endodermal lineage cells, where red indicates higher differentiation potential states and blue indicates more differentiated states. (**D**) UMAP plot of endodermal lineage cells colored by developmental stages of trj and mj. (**E**) Dot plot showing expression levels of representative marker genes distinguishing the two major endodermal lineages. Dot color reflects average expression within each cluster, and dot size indicates the proportion expressing cells. (**F**) Pseudotime trajectory analysis visualized on the UMAP plot. Cells are colored by pseudotime score. (**G**) RNA velocity streamlines overlaid on the UMAP plot, indicating the predicted directions of cell state transition within the endodermal lineage. (**H**) Schematic illustration summarizing the two major endodermal lineages and their respective differentiation fates. (**I**) (Top) Heatmap showing cross-species transcriptomic similarity between ascidian and mouse endodermal lineages. AUROC scores were calculated using expression of one-to-one orthologous genes. (Bottom) Schematic diagram of mouse definitive endoderm and visceral endoderm lineages. PrE, primitive endoderm; ParE, parietal endoderm; VE, visceral endoderm; emVE, embryonic visceral endoderm; exVE, extra-embryonic visceral endoderm; TE, trophectoderm; ExE, extra-embryonic ectoderm; EPI, epiblast; DE, definitive endoderm.

To reveal the difference between them, the gene expression profiles were exhibited (Figure 3E). The trunk endodermal marker genes *Lhx3* and *Nkx2-1* (*TTF1*) were expressed in the high differentiation potential endoderm 1 and endoderm 2. *Foxp4, Otx1* expressed in the high differentiation potential endoderm 1 and *Sox7, FGF16* expressed in the high differentiation potential endoderm *2*. Hox10, the marker of the endodermal strand cells in larval tail,^23^ was detected in low differentiation potential endoderm 1 and 2. Meanwhile, *Amyloid precursor-like protein* (*APLP*) and *Carboxypeptidase Z* (*CBPZ*) were detected in the low differentiation potential endoderm 1; *CBPE, Six1, Low-density lipoprotein receptor 1* (*LRP1*) and *LRP5* were expressed in the low differentiation potential endoderm 2 (Table S2). The results indicated that the high differentiation potential endoderm cells derived from endoderm in larval trunk and the low differentiation potential endoderm derived from the endodermal strand in larval tail.

In order to further reveal the cell lineage relationships, pseudotime trajectory of these cells were constructed using Monocle3. We designated the high differentiation potential endoderm 1 population, which exhibited the highest differentiation potential, as root in pseudotime ordering, and ordered all cell types of endoderm lineage. The results showed that one branch developed from high differentiation potential endoderm clusters to the stomach progenitor clusters; and the other one was from the low differentiation potential endoderm clusters to the intestine progenitor clusters (Figure 3F). RNA velocity analysis also indicated that stomach progenitor clusters, endostyle cluster, and immune cluster originated from the high differentiation potential endoderm cells, while intestine progenitor clusters originated from the low differentiation potential endoderm cells (Figure 3G). Combining the endodermal cell lineages and pseudo-time analysis results, we inferred that high differentiation potential endoderm cells are corresponded to trunk endoderm lineage which could generate the stomach progenitors, while low differentiation potential endoderm cells are corresponded to endoderm strand lineage which could generate the intestine progenitors (Figure 3H).

To determine the evolutionary relationships of the endodermal cells between ascidian and vertebrates, we compared ascidian endodermal cells at trj stage with mouse embryonic endoderm cell types from E4.5 to E8.75 stage.^6^ The results showed that the ascidian high differentiation potential endoderm cell types 1 and 2 are both corresponded to the mouse definitive endoderm (DE) lineages (including epiblast) and ectoderm lineages (including trophectoderm and extra-embryonic ectoderm). While the low differentiation potential endoderm cell types 1 and 2 are both correspond to the visceral endoderm (VE) lineages including primitive endoderm, embryonic visceral endoderm, extra-embryonic visceral endoderm and parietal endoderm (Figure 3I).

### The identification of stomach and intestine progenitor cells

To investigate the potential differentiation signals of stomach and intestine progenitor cells, we selected the digestive tract related cells from mj stage and further divided them into sub-types. Seven sub-types were identified based on maker genes, including three sub-types of stomach progenitors (aminopeptidase N*+,*), three sub-types of intestine progenitors (*Hox10+*), and a group of cells expressing stem-cell markers such as *Piwi, Tolloid, Six3/6* (labeled as digestive progenitors) (Figure 4A, Table S3). To verify the cell type annotation, we compared our scRNA-seq data with the published bulk RNA-seq data of digestive tract in ascidian *Ciona savignyi*.^24^ The results showed that stomach progenitors of *S. clava* had the highest expression similarity with the stomach and anterior intestine of *C. savignyi,* while the intestine progenitors had the highest expression similarity with the middle intestine and posterior intestine (Figure S3).

**Figure 4.**
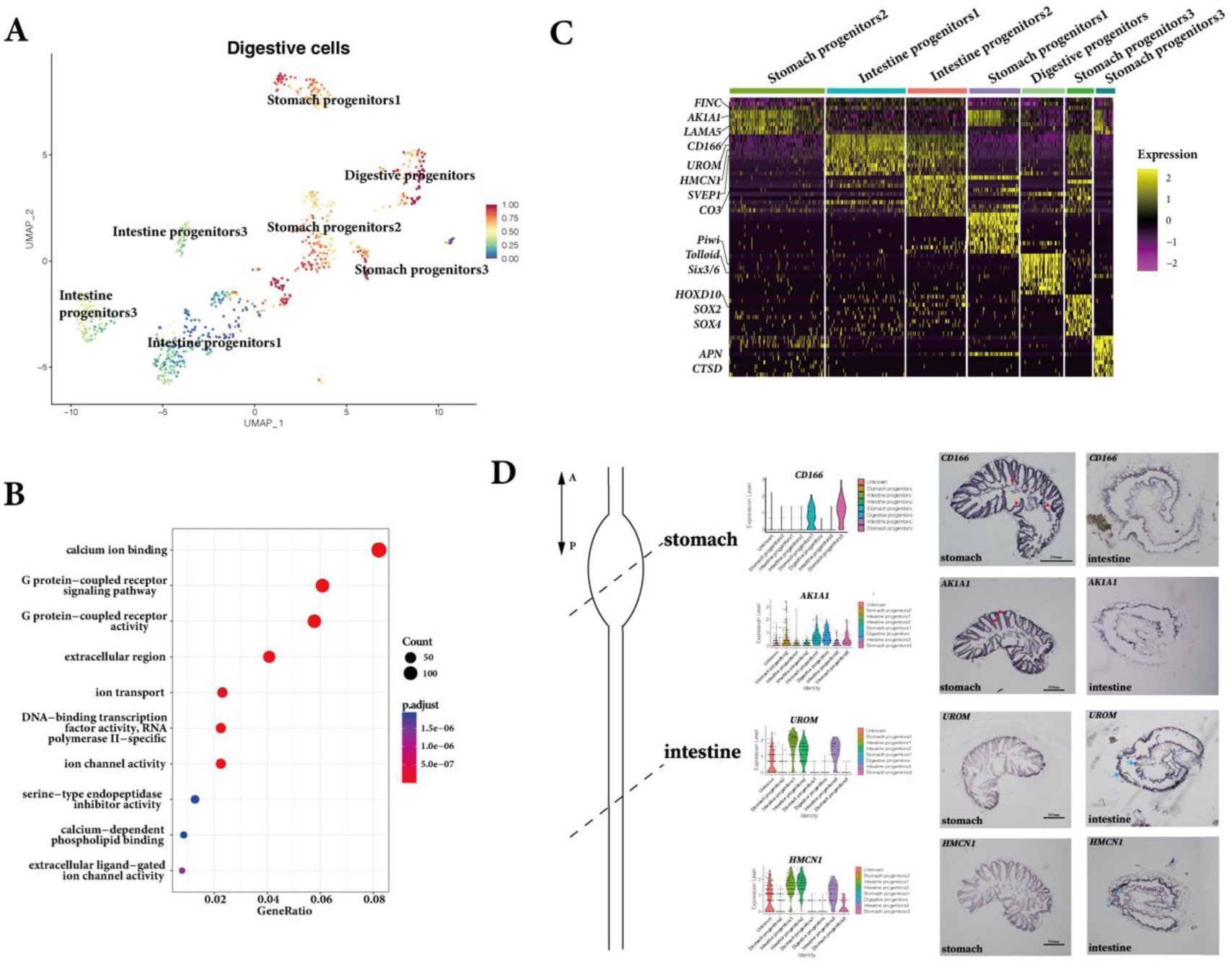
Characterization of stomach and intestine progenitor cells. (**A**) UMAP plot showing sub-clustering of digestive tract cells at the metamorphic juvenile stage. Cells are colored according to CytoTRACE-inferred differentiation scores, with red indicating higher differentiation potential. (**B**) Dot plot showing enriched Gene Ontology (GO) terms for the top 100 differentially expressed genes in digestive cells compared to all other cells at a certain stage. (**C**) Heatmap displaying the expression of representative marker genes in the major digestive tract cell types. Each column represents a cell type, and each row represents a marker gene. (**D**) Schematic diagram of the adult digestive tract (stomach and intestine), with dotted lines indicating the anterior-posterior levels of transverse sections (left). Violin plots showing the expression levels of marker genes (CD166 and AK1A1) in stomach progenitor cells and maker genes (UROM and HMCN1) in intestinal progenitor cells (middle). *In situ* hybridization results validating marker gene expression in ascidian digestive tract (right). Red and blue arrows indicate signals in stomach and intestinal regions, respectively.

To identify the biological function of these seven sub-types, the top100 differentially expressed genes (DEGs) were identified by comparing these cells agaist all other cells at mj stage for Gene Ontology (GO) analysis. The result showed that the genes of digestive cell types were enriched in calcium ion binding, G protein-coupled receptor activity, G protein-coupled receptor signaling pathway, extracellular matrix region, ion transport and DNA-binding transcription factor activity (Figure 4B). The gene expression analysis showed that extracellular matrix and digestive enzymes genes such as *FINC, CD166, LAMA5, AK1A1, APN, CTSD* were preferentially expressed in stomach progenitors. While, *HPGDs, UROM, HMCN1, SVEP1, SOX2, Hox10* were expressed in intestine progenitors (Figure 4C, Table S3). The expression patterns of these markers were experimentally verified in digestive tract of adult ascidians by *in situ* hybridization. The signals of *CD166* and *AK1A1* (stomach progenitor markers) were mainly expressed in the folds of adult stomach, while *UROM* and *HMCN1* (intestine progenitor markers) showed stronger signals in the inner fold of intestinal epithelium (Figure 4D). The expression pattern of marker genes further supported the contribution of these digestive progenitors to the digestive organs in adult.

### Cell differentiation signals of stomach and intestine progenitors

To investigate the cell differentiation of stomach and intestine lineages, a pseudo-time developmental trajectory of digestive cells at mj stage was constructed using Monocle2. The trajectory showed two distinct directions, with one differentiating into stomach progenitors and the other into intestine progenitors (Figure 5A, Figure S4A). The branched gene expression analysis on the distinct directions of stomach and intestine progenitors highlighted the genes on each direction (Figure 5B, Table S4). *MASP1*, *ITLN2* and *LARP1* were highly expressed in stomach direction. *KLF7, LRP4* and *Notch2* were highly expressed in intestine direction. *FOS* was differently expressed in the two branches (Figure 5B).

**Figure 5.**
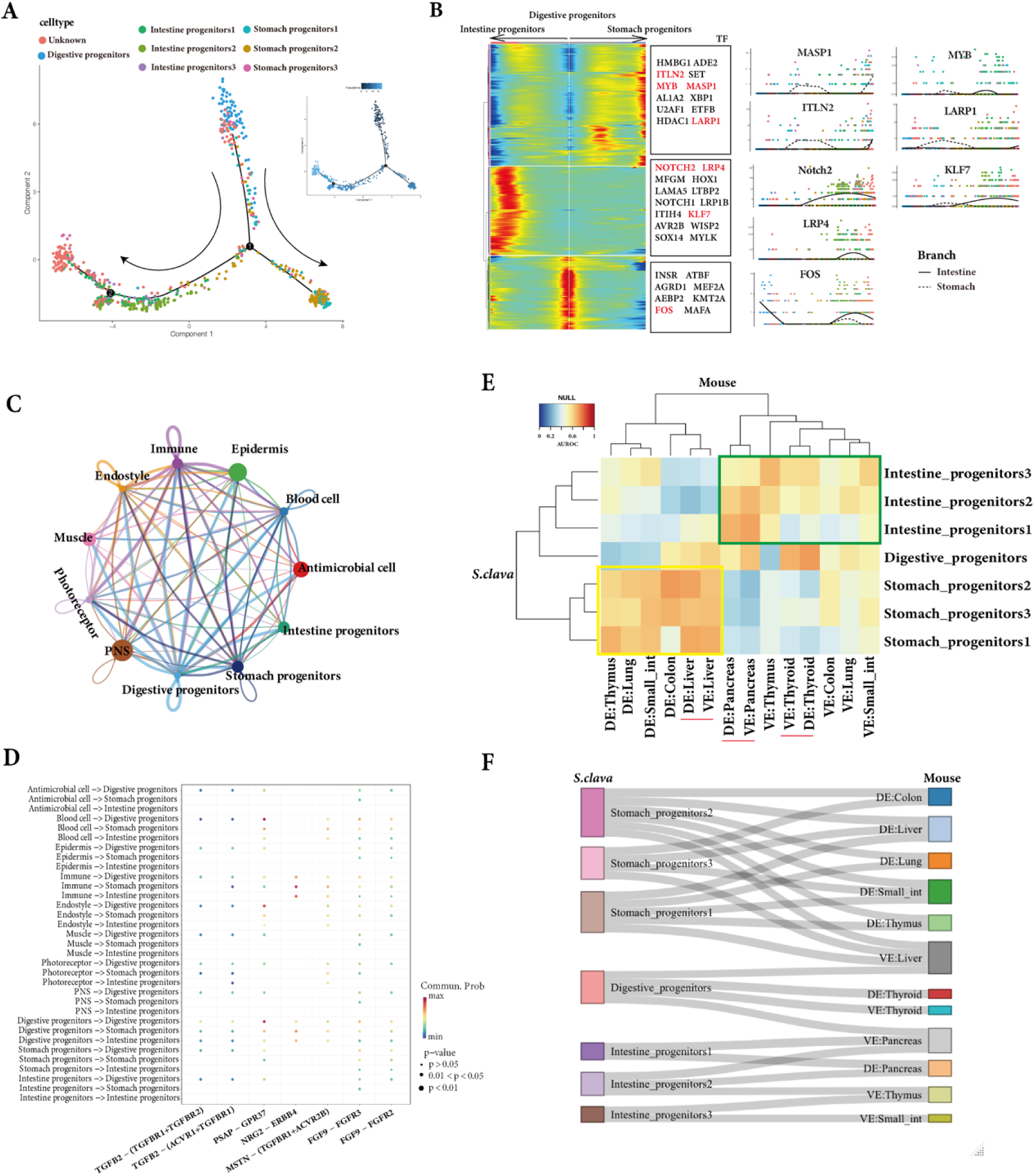
Lineage-specific signaling and cross-species comparison of stomach and intestine progenitor cells. (**A**) Developmental trajectory of digestive progenitor cells, colored by digestive cell types. Arrowheads indicate inferred directions of differentiation. The inset (right) shows the same trajectory colored by pseudo-time scores, where darker shades indicate closer to the root cells. (**B**) Heatmap showing gene expressions along the differentiation trajectories of intestinal and stomach progenitors. Lineage-specific transcription factors (TFs) are labeled. The expression dynamics of representative TFs (red) are displayed along the corresponding trajectories. (**C**) Network showing inferred signaling interactions between digestive progenitor cells and other cell types at the same developmental stage. Edge thickness reflects the strength of inferred communication. (**D**) Dot plot showing the expression levels of ligand–receptor pairs between digestive tract cell types and interacting cell types. Dot color indicates communication probability, while dot size represents statistical significance (p-value from one-sided permutation test). (**E**) Heatmap of cross-species transcriptomic similarity between ascidian and mouse digestive progenitor cells. AUROC scores were calculated based on the expression of one-to-one orthologous gene pairs. (**F**) Sankey diagram showing homologous relationships between ascidian and mouse digestive tract progenitor cells (AUROC > 0.7). Small_int, small intestine.

To construct the cell-cell communication during the development of the digestive tract, the CellChat analysis of all cell types to digestive-related cells at mj stage were conducted (Figure 5C). The results showed that the interactions mediated by ligands TGFB2 and FGF9 present in digestive related cell types, which belong to the *TGF-β* signaling pathway and *FGF* signaling pathway respectively (Figure 5D). Furthermore, the expression level of *FGF* signaling was higher in digestive progenitors than in stomach and intestine progenitors, and TGF-β signaling was higher in digestive progenitors and stomach progenitors than in intestinal progenitors (Figure S4B). Meanwhile, PSAP-GPR37 and NRG2-ERBB4 interaction signal were also found in cell-cell interaction of digestive related cells (Figure 5D).

To explore the homologous relationship of the digestive progenitors between ascidian and vertebrate, we compared the expression feature of digestive progenitor cells of ascidian with mouse digestive organ progenitors (Table S5). The stomach progenitors 1, 2 and 3 from ascidian exhibited higher similarity with DE-derived progenitors in mouse including DE cells in thymus, lung, liver, small intestine and colon. The intestine progenitor 1 exhibited higher similarity with both DE and VE progenitors from pancreas in mouse, and the intestine progenitor 2 and 3 exhibited higher similarity with VE-derived progenitors in mouse, including VE cells in thymus, thyroid, lung, pancreas, small intestine and colon (Figure 5E and 5F), which indicated a different cell lineage composition of digestive organs between ascidian and mouse.

## Discussion

In ascidian, there is a unique retrogressive metamorphosis process.^25,26^ Due to the unique lifestyle strategy of ascidian, the development processes of endoderm-derived organs are unsynchronized with ectoderm and mesoderm derived organs, which differs from most vertebrates.^27–29^ This allows us to better trace the formation of digestive tract. In this study, we present an extensive analysis of single-cell transcriptomic of ascidian *S. clava* from five developmental stages that cover the metamorphosis process.

### Ascidian digestive tract progenitor cells originate from different endodermal cell populations with distinct differentiation potentials

In ascidians, the endodermal cells of the tadpole larva are distributed in the trunk and tail, corresponding to the trunk endoderm and tail endodermal strand, respectively.^28,30^ The digestive tract develops during metamorphosis from these two primordial tissues. The endodermal cells undergo epithelialization to form a tubular structure from stage 27 (tail regressing larva stage); and the development of the digestive tract is completed at stage 37 (metamorphic juvenile stage) after body axis rotation.^31^ According to previous cell tracing experiment, the anterior and posterior parts of the digestive tract are formed through distinct processes from the posterior endoderm and the endodermal strand, respectively. However, it still has been largely unknown about the characteristics of these primordial cells. Here, we identified high and low differentiation potential endoderm cells at trj stage. The further pseudo-time trajectories analyses inferred that the high differentiation potential endoderm cells generated into the stomach progenitor, and the low differentiation potential endoderm cells generated into the intestine progenitor. These results revealed the different developmental origins of stomach and intestine organs and a significant difference in differentiation potentials between two groups of endoderm cells. These distinct differentiation potentials may have functional consequences for the following organ specialization, which also could help to explain different regions of the digestive tract exhibit distinct cellular compositions and regenerative capacities.^32,33^

### Conserved molecular signaling in digestive tract regionalization between ascidians and vertebrates

During organ formation, cell identity and the tissue morphogenesis must be tightly coordinated. Several signaling pathways, including those of FGF, Wnt, hedgehog, BMP and retinoic acid are used reiteratively during endoderm organ development in vertebrates.^34–37^ Among them, FGF signaling plays a particularly pivotal role in the early regionalization of the gut tube, where it is required for establishing distinct domains along the anterior-posterior (A-P) axis.^38^ The cell-cell communication analysis of ascidians also identified the FGF signaling among ascidian digestive progenitor cells. This observation suggests evolutionary conservation of FGF-mediated patterning mechanisms in gut development, highlights the conserved molecular but not cellular logic underlying digestive tract regionalization across chordate evolution.

### Dual origins of digestive tract progenitor cells in both ascidians and mammals

Recent study demonstrates both embryonic and extraembryonic cells contribute to the emerging gut endoderm in mouse.^39^ Our work also showed dual origins of digestive tract progenitors, with some cell populations corresponded to the mouse DE lineages (embryonic cells), the other cell populations correspond to the VE lineages (extraembryonic cells). The lineage tracing experiments in sea urchin^40,41^ also show that digestive tract is derived from the endoderm, but the foregut (together with the oral midgut) and the hindgut (together with the aboral midgut) are derived from different macromere lineages. These indicated conserved dual origins of digestive tract progenitor cells in deuterostomes.

Nevertheless, the cells contribute to the landscape of the specialized endodermal organs seems different between ascidians and mammals. The stomach progenitor cells and intestine progenitor cells in ascidians corresponded to the DE and VE lineages that constitute the digestive organs, rather than corresponding to specific organs in mouse. This suggests that the cell fates of ascidian stomach and intestine progenitors restricted earlier, which is distinct from vertebrates. Moreover, recent study showed the extraembryonic gut cells (VE lineage) of mouse undergo programmed cell death and neighboring embryonic cells clear their remnants via non-professional phagocytosis, so that VE lineage cells are no longer detected in the wild-type gut.^42^ These indicated a different cell fate determination and contribution to the digestive tract between ascidians and mammals.

Overall, the dual lineage origin of the digestive tract appears to be a fundamental feature established during development in deuterostome. The complex digestive tract development observed in vertebrates may represent an extension and elaboration of the cellular heterogeneity already present in the endodermal cells of chordate ancestors.

## Conclusions

This study provided the single-cell atlas during metamorphosis of ascidians, and elucidated the pseudo-trajectories of endoderm differentiation in *S. clava*. Our work revealed that the digestive tract originated from two endodermal progenitor populations with distinct differentiation potentials in ascidians. Comparative analyses with vertebrates support a shared dual origin but divergent differentiation trajectories of digestive tract lineages. Together, these findings provide a cellular framework for understanding digestive tract specialization and the evolutionary diversification of endodermal lineages across chordates.

## Materials and Methods

### Animals and sample collection

Adult *S. clava* were obtained from Xunshan Company (Weihai, China) and temporarily maintained in a recirculating aquarium system at 18 °C in the laboratory, with continuous aeration provided by air pumps. Healthy adults exhibiting normal gamete release were selected. Sperm and eggs were collected separately and mixed for fertilization, which was allowed to proceed for 30 minutes before transferring the zygotes to culture dishes. The culture dishes were subsequently incubated at 18 °C for further development. Embryos were monitored periodically according to the established developmental staging for ascidians.^43^ At the desired developmental stages, larvae or juveniles were collected for subsequent experiments. Samples were collected at five representative developmental stages: hatched swimming larva, tail-regressing larva, tail-regressed larva, metamorphic juvenile, and juvenile stages.

### Single-cell isolation and scRNA-seq library construction

Samples from five developmental stages were used for single-cell isolation and library construction. Three biological replicates were prepared. For scRNA-seq library preparation, gel beads in emulsion (GEMs) were generated using the Chromium Controller (10x Genomics). Libraries were constructed using the Chromium Next GEM Single Cell 3’ Reagent Kits v3.1 (10x Genomics), following the manufacturer’s protocol. The resulting libraries were sequenced on the Illumina NovaSeq platform.

### Data analysis for scRNA-seq

For the scRNA-seq data generated using the 10x Genomics Chromium platform, each of the biological replicate was processed independently. Raw sequencing data were processed using Cell Ranger to generate gene-barcode matrices for each sample, employing the default parameters. A previously published genome assembly was used as the reference.^15^ Gene expression matrices were then imported into Seurat v4.0 to create Seurat objects.^44^ Standard preprocessing steps including quality filtering, normalization, integration, and clustering were performed. Cells were initially filtered based on expression metrics, retaining those with nCount between 0 and 30,000 and nFeature between 300 and 1,000. Cells exhibiting high mitochondrial gene content (defined as >15% of total counts) were excluded. High-quality datasets were normalized and scaled using SCTransform, followed by dimensionality reduction. The biological replicates were then integrated using the IntegrateData function to remove batch effects and enable joint analysis. The integrated dataset was subjected to principal component analysis (PCA), and unsupervised clustering was performed using the shared nearest neighbor algorithm with a resolution of 0.5. Marker genes for each cluster were identified using the FindAllMarkers function.

### Cell type annotation

For both the single-stage and integrated single-cell datasets covering five developmental stages, differentially expressed genes (DEGs) were identified using the FindAllMarkers function in Seurat with default parameters. For each dataset, the top 30 DEGs were selected based on fold change and statistical significance (adjusted p-value), and their functional annotations were retrieved from the Swiss-Prot database (https://www.ebi.ac.uk/uniprot/index). Given that our dataset spans five distinct stages of ascidian metamorphosis, we performed cell clustering and annotation independently for each stage. Cell types or cluster identities were manually assigned based on the expression of cluster-enriched marker genes in each stage-specific single-cell dataset. Final cell type annotations were consolidated based on the clustering results from the integrated dataset, allowing for consistent comparison across stages.

### GO analysis

GO analysis was performed with the package clusterProfiler4.0.^45^ Only biological process GO terms with a *p*-value < 0.05 were selected. The results were further visualized with the ggplot2 package.

### RNA velocity analysis

Loom files containing unspliced and spliced reads were generated from velocyto.py for downstream analysis with default parameters. The python package scVelo ^46^ was applied to estimate RNA velocity with a dynamic model by default parameters. The RNA velocities were projected into UMAP embedding and visualized by cell types.

### Cell trajectory analysis of the endodermal lineage cells and digestive progenitors

Pseudotime analysis of endodermal lineage cells was performed using the Monocle3 package.^47^ The gene expression matrix and UMAP coordinates were imported into a CellDataSet object. Cell trajectories were constructed using the learn_graph function with the parameter minimal_branch_len = 3. To infer pseudotime, the order_cells function was applied, and root cells were manually selected to anchor the trajectory. For digestive progenitor cells, pseudotime analysis was conducted using Monocle2.^47^ Genes highly expressed in these clusters (log_avg2FC > 0.25) were identified using the differentialGeneTest function to detect dynamic gene expression changes along pseudotime. To analyze branch-specific expression patterns, the BEAM function was applied with branch_point = 1. Genes exhibiting branch-dependent expression were exported, and transcription factors were selectively filtered from this list. The plot_genes_branched_pseudotime function was then used to visualize temporal expression patterns of these TFs along divergent trajectories.

### Cell-cell interaction analysis

Cell‒cell communication analysis was performed using the CellChat package.^49^ The analysis utilized the CellChatDB.human database, which includes interactions from secreted signaling, extracellular matrix–receptor binding, and direct cell–cell contact ligand–receptor pairs. The resulting Seurat object was converted into a CellChat object and processed according to the recommended workflow in the CellChat manual.

### Comparison across species between ascidian and vertebrates

Single-cell RNA-seq data for mouse endodermal tissues were obtained from the GEO database (accession number: GSE123046). Cross-species comparison was performed based on one-to-one orthologous genes between *S. clava* and mouse. The MetaNeighbor package^50^ was used to assess cell-type similarity across species based on gene expression profiles. Sankey diagrams were generated using the networkD3 package (https://christophergandrud.github.io/networkD3/) to visualize correspondence between cell clusters in *S. clava* and mouse.

### *In situ* hybridization experiment

The digestive tract of healthy individuals was carefully dissected using sterilized instruments. After isolating the entire digestive tract, the stomach was separated from the intestinal region based on distinct external morphological features. Fresh samples were fixed in 4% paraformaldehyde in artificial seawater at 4 °C overnight. Subsequently, the tissues were cryoprotected by immersion in 30% sucrose in PBS. The tissues were embedded in O.C.T. compound (Sakura 4583) and snap-frozen in liquid nitrogen. Cryosections were prepared using a Leica CM3050 cryostat and mounted on adhesive microscope slides. Digoxigenin (DIG)-labeled RNA probes were designed (Table S6) and synthesized by in vitro transcription using SP6 and T7 RNA polymerases (Thermo Fisher, EP0131 and EP0111). *In situ* hybridization (ISH) was performed in a humidified dark chamber to prevent tissue desiccation. Tissue sections were permeabilized with 10 μg/mL Proteinase K for 30 minutes at 37 °C, followed by washing in PBST (0.1% Tween-20 in PBS). Sections were then post-fixed with 4% PFA in PBS for 30 minutes at room temperature, followed by additional PBST washes. For hybridization, tissues were first incubated in pre-hybridization buffer (50% formamide, 5× SSC, 0.1% Tween-20) for 10 minutes at room temperature, and then transferred to hybridization buffer (50% formamide, 5× SSC, 0.1% Tween-20, 5× Denhardt’s solution, 100 μg/mL yeast tRNA, 50 μg/mL heparin) at the hybridization temperature for 4 hours. Fresh hybridization buffer containing RNA probes (0.5 ng/μL) was then applied, and the tissues were incubated for 20 hours at the same temperature. Post-hybridization, sections were washed four times in hybridization wash buffer (50% formamide, 5× SSC, 0.1% Tween-20), followed by four washes in PBST. Slides were then incubated overnight at 4 °C in 1% blocking buffer (Roche) containing the anti-DIG antibody. Finally, chromogenic detection was performed using NBT/BCIP substrate.

### Statistics analysis

All the statistical analyses were performed using R (version 3.6). Student’s *t*-test and Spearman’s rank correlation analysis were utilized in this study.

## Availability of data

All data generated or analyzed during this study are included in the manuscript and supporting files. The RNA-sequencing datasets generated in this study have been deposited at NCBI under the BioProject accession number PRJNA1122959.

## Competing interests

The authors declare no competing interests.

## Acknowledgments

The authors thank Professor Sebastian Shimeld (Department of Zoology, University of Oxford) for constructive comments and suggestions on the manuscript.

## Funding

This work was funded by the National Natural Science Foundation of China (Grant No. 32522017, 32370461), the Science & Technology Innovation Project of Laoshan Laboratory (No. LSKJ202203001, LSKJ202203002), the Natural Science Foundation of Shandong Province (ZR2023MD029), and the Taishan Scholar Program of Shandong Province, China (B.D.).

## Authors’ contributions

Conceptualization: B.D., J.W., and H.Y. Project supervision: B.D., and J.W. scRNA-seq library generation: Y.G., W. Z., J.W., Data analyze: Y.G., and P.L. Technical and experimental support: Y.G., J.B., and H.Y.. Writing—original draft: Y.G. Writing—review and editing: J.W., and B.D.

## Notes

### Competing Interest Statement

The authors have declared no competing interest.

